# Synaptic dynamics govern spatial integration in mouse visual cortex

**DOI:** 10.1101/2025.05.11.653343

**Authors:** Jennifer Y. Li, Celine M. Cammarata, Lindsey L. Glickfeld

## Abstract

Neurons in primary visual cortex are often suppressed by stimuli extending beyond their receptive fields. This surround suppression is proposed to reduce the redundancy of encoding large stimuli and support scene segmentation. We find that surround suppression decreases firing rates in mouse primary visual cortex by accelerating the decay of visually-evoked responses and reducing response duration. The rapid decay of visual responses at large sizes is enhanced by increased contrast, reduced by locomotion, and invariant to stimulus orientation, consistent with the engagement of a network mechanism. While fast-spiking interneurons have faster dynamics relative to neighboring pyramidal cells, the dynamics of somatostatin-expressing interneurons are delayed. At the subthreshold level, the rapid decay of visual responses with increasing size is due to a delayed removal of both synaptic excitation and inhibition below baseline levels following visual input. We propose that the delayed activation of somatostatin-expressing interneurons drives a network-wide suppression and accelerates the decay of the visual response. Thus, these data identify a key role for synaptic network dynamics in regulating both spatial and temporal integration in mouse visual cortex.

## Introduction

The spatial context of a visual stimulus robustly impacts sensory processing and perception^1–3^. Stimuli that effectively fill the classical receptive field (CRF) of a neuron drive stronger responses than those that extend beyond the boundaries of the CRF^4–6^. This nonlinear integration, known as surround suppression, sparsens sensory responses and is thought to reduce the redundancy of encoding^7–9^. It also drives a decrease in the perceived contrast and size of larger stimuli^10–12^.

The phenomenon of surround suppression is present at all stages of the visual processing hierarchy^2^, with important distinctions. In the retina, the effects of surround suppression extend only a few degrees beyond the CRF^13^, whereas neurons in the primary visual cortex (V1) are sensitive to much larger surrounds^14–16^. Thus, while some of the effects of surround suppression in V1 are inherited along the feedforward pathway, a substantial portion is thought to be locally generated through the recruitment of lateral and top-down inhibitory circuits^17–19^. Indeed, in the cortex, surround suppression has a relatively late onset^20,21^, consistent with the delayed recruitment of cortical inhibition. This delayed onset of surround suppression suggests that it also plays a role in temporal integration in addition to spatial integration. This is supported by observations that increasing stimulus size decrease the time course of perceptual integration^22–25^. However, it has remained unclear how inhibitory mechanisms in the cortex link this response modulation across spatial and temporal domains.

Multiple models have been proposed for inhibitory control of spatial integration. In one model, activation of the surround drives an increase in inhibition which decreases the ratio of excitation to inhibition, thereby decreasing visual responses^10,26^. An alternative model, in which the cortex exists in an inhibition stabilized network^27,28^ (ISN), argues that transient inhibition subsequently drives a decrease in recurrent excitation which ultimately results in a decrease of both excitation and inhibition. Intracellular recordings that directly measure excitation and inhibition during visual stimulus presentation provide disparate evidence for each of these models^9,17,29–31^. However, these studies make their measurements in the steady-state condition and in response to sustained stimuli, which given the dynamic nature of network stabilization, may obscure key events.

To investigate the inhibitory mechanisms underlying surround suppression, and its effects on temporal integration in V1, we made electrophysiological recordings in response to brief, static stimuli varying in size. We find that increasing stimulus size robustly reduces the average firing rate of V1 neurons while leaving the peak firing rate largely unchanged, suggesting a major effect of surround suppression on the dynamics of visual responses. Indeed, large stimuli accelerate of the decay of visual responses, driving activity below spontaneous firing rates, and decreasing response duration. The decrease in response duration is enhanced by contrast, insensitive to orientation, and reduced by locomotion, consistent with the engagement of network mechanisms. Intracellular recordings reveal that the decrease in response duration is due to a speeding of the dynamics of the underlying excitatory and inhibitory synaptic currents. After a transient increase in excitation and inhibition, both are suppressed below spontaneous levels, with little change in the balance of excitation and inhibition. Thus, our data provide support for the role of inhibition stabilization in surround suppression, and a mechanistic link between temporal and spatial integration in the cortex.

## Results

### Surround suppression is generated by increasingly transient visual responses

To systematically measure the dynamics of visual responses while engaging different amounts of surround suppression, we made *in vivo* extracellular recordings of neurons across all layers of primary visual cortex (V1) in awake, head-fixed mice while presenting static grating stimuli (0.1 s duration; contrast: 80%; spatial frequency: 0.1 cycles per degree) of varying diameter (diameter: 7.5°-120°; **Figure 1a-b**). To ensure receptive fields were well-matched to the visual stimulus, only units that significantly increased firing rate at the smallest size were included. Selecting units with receptive field centers within 10 degrees of the stimulus center yielded similar results (**Figure S1a-b**).

**Figure 1.**
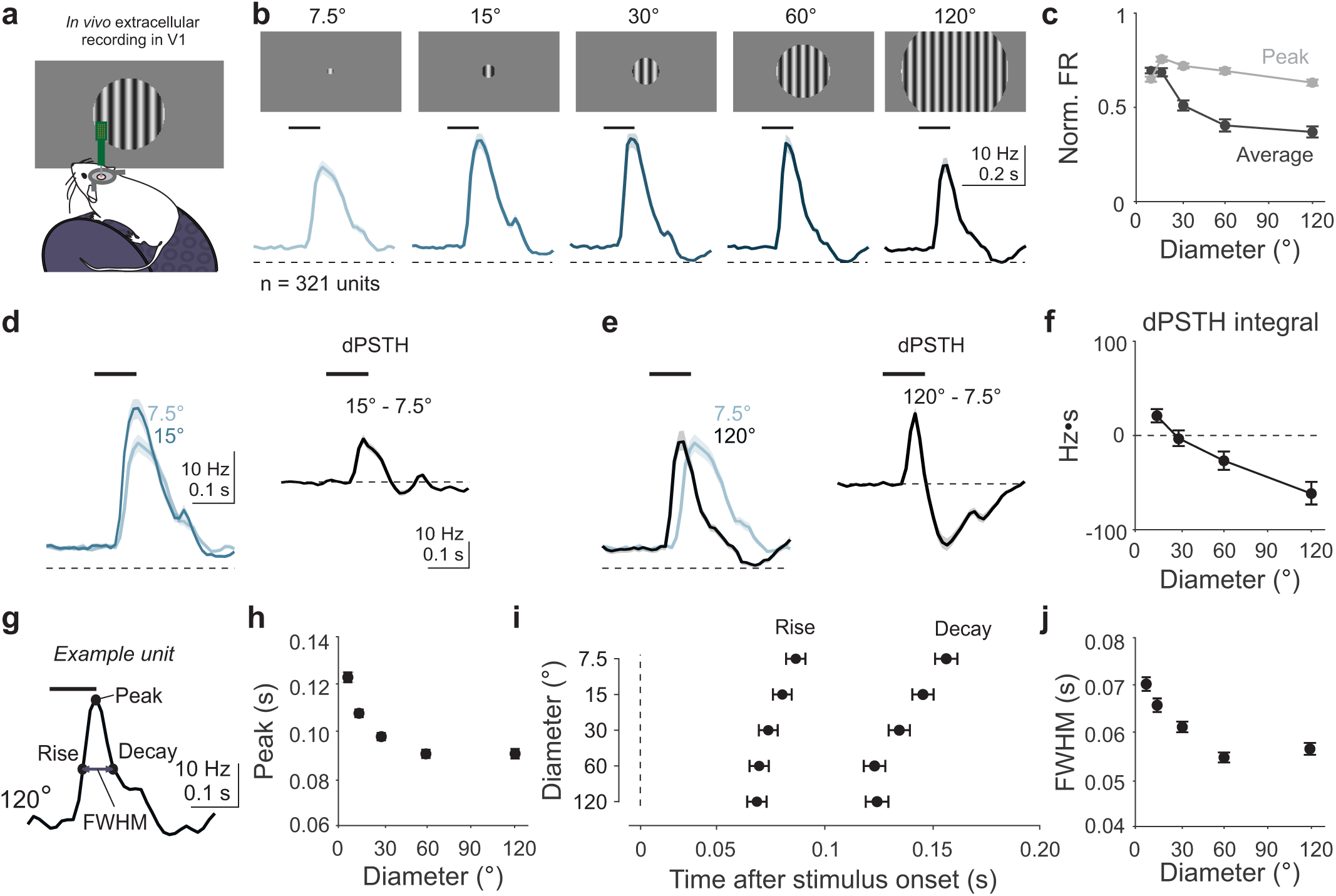
Surround suppression is mediated by a decrease in response duration. (a) Schematic of *in vivo* extracellular recordings performed in head-fixed mice. (b) Average visually-evoked post-stimulus time histogram (PSTH) in response to presentation of static gratings of different diameters (n = 321 units). Shaded error is SEM across units. Horizontal dashed line is 0 Hz; thick horizontal line is time of stimulus. (c) Average normalized firing rate as a function of stimulus diameter measured using average (dark grey) and peak (light grey) of the PSTH. Error is SEM across units. (d) Left, comparison of PSTHs for 7.5° (light blue) and 15° (dark blue) stimuli; right, pointwise subtraction of stimuli on left are used to create the difference PSTH (dPSTH). (e) Same as (d) but for comparison with responses to 120° stimulus. (f) Integral of the dPSTH as a function of stimulus diameter. (g) Illustration of 50% rise, 50% decay, peak and full width at half-max (FWHM) for PSTHs from an example unit in response to 120° stimulus. (h) Time to peak relative to stimulus onset across different diameters, averaged across responsive units. (i) Same as (h), for time to 50% rise and decay. (j) Same as (h), for FWHM. Only units responsive to each stimulus condition are included: n = 321, 289, 275, 252, 248.

Consistent with previous findings, average visually-evoked responses (0-0.35 s after stimulus onset) undergo robust surround suppression^4,6,32^ (suppression index (SI)-0.63 ± 0.03; n = 321 cells, 15 mice; Student’s t-test, p = 8.41e-62; **Figure 1c**). However, the peak response is more weakly modulated with increasing stimulus size (SI-0.36±0.02; paired t-test, average vs peak: p = 5.13e-30). The stronger dependence of the average, compared to the peak, visually-evoked response on size is consistent across all cortical layers (**Figure S1c-e**). Inspection of the peri-stimulus time histograms (PSTHs) reveals that the divergence of peak and average response is due to differences in the decay of the visual response: when the stimulus is small, neurons undergo a transient increase in firing rate before decaying nearly monotonically back to baseline; at large sizes, the decay of the PSTH transiently undershoots the baseline firing rate before returning to baseline (**Figure 1b**).

To visualize the change in temporal dynamics across stimulus size, we calculated the difference PSTH (dPSTH): the pointwise difference of each unit’s PSTH in response to the smallest size (7.5°) from its PSTH to increasing sizes (**Figure 1d-e**). The dPSTH for a 15-degree stimulus has a single positive peak that reflects the decreased latency and increased amplitude of the response compared to the 7.5° stimulus (**Figure 1d**). With increasing size, the dPSTH exhibits a biphasic time course with an early positive peak followed by a longer-lasting negativity (**Figure 1e**). These dynamics reflect changes in both the timing and number of spikes. The time to peak of the positive transient in the dPSTH is significantly reduced with increasing size (one-way ANOVA, p = 0.02), consistent with decreased latency in the visual response. In addition, the integral of the dPSTH becomes increasingly negative, reflecting a decrease in the total number of spikes with larger sizes, especially during the late portion of the visual response (one-way ANOVA, p = 5.20e-14; **Figure 1f**). This biphasic dPSTH is the product of intrinsic cortical dynamics, not an artifact of the stimulus duration, as a 1 s stimulus evokes similar dynamics (**Figure S1f-i**). Thus, the dPSTH reveals that increasing size decreases both the latency and number of visually evoked spikes.

We further quantified the effect of stimulus size on visual response dynamics by measuring the time to rise, decay, peak and duration (full-width at half max, FWHM) of the response (**Figure 1g**). Consistent with our measure of the dPSTH, we find that increasing size shifts visual responses to be earlier, significantly reducing the time to peak (one-way ANOVA, p = 4.54e-44; **Figure 1h**). Underlying this shift is a decreased latency of time to 50% rise (one-way ANOVA, p = 2.10e-64, **Figure 1i**), and decay (p = 5.08e-67). Notably, there is a larger effect of size on decay than on rise (change in rise from 7.5° to 120°: −20.5 ± 1.3 ms, change in decay: −39.4 ± 2.5 ms; change in rise versus decay: p = 4.02e-18, paired t-test). Consequently, we find that increasing size results in a significant decrease in FWHM (one-way ANOVA, p = 5.13e-22; **Figure 1j**), indicating that responses become more transient. Therefore, stimulus size strongly modulates the dynamics of visual responses, reducing response latency and duration, which ultimately decreases average firing rates and manifests as surround suppression.

### Decrease in response duration is selective to increases in stimulus size

Surround suppression is recruited more robustly with increasing contrast^33^. To determine how increasing contrast impacts the size-dependent dynamics in visual cortex, we presented an expanded stimulus set of the same range of sizes (7.5°-120°) at four different contrasts (0.1, 0.2, 0.4, 0.8; **Figure 2a-b**). We find that increasing stimulus contrast enhances surround suppression in a manner similar to previous studies^35,36^, but only when measuring activity with the average FR (two-way ANOVA, interaction of size and contrast, p = 2.04e-08; **Figure S2a**), not the peak (p = 0.10; **Figure S2b**). Consistent with our hypothesis that surround suppression is mediated by size-dependent acceleration of visual response dynamics, increasing contrast enhances the effect of size on the time to peak (two-way ANOVA, interaction of size and contrast, p = 0.03; **Figure S2c**). Increasing contrast also significantly reduces the time to rise (two-way ANOVA, effect of contrast: p = 1.03e-94; **Figure 2c**), decay (p = 1.52e-141), and FWHM (p = 2.39e-46). However, unlike the time to peak, we find no significant interaction of contrast and size for rise (two-way ANOVA interaction of size and contrast, p = 0.34), decay (p = 0.09), and FWHM (p = 0.28), indicating that increasing contrast significantly enhances these measures, but does not change their size-dependence.

**Figure 2.**
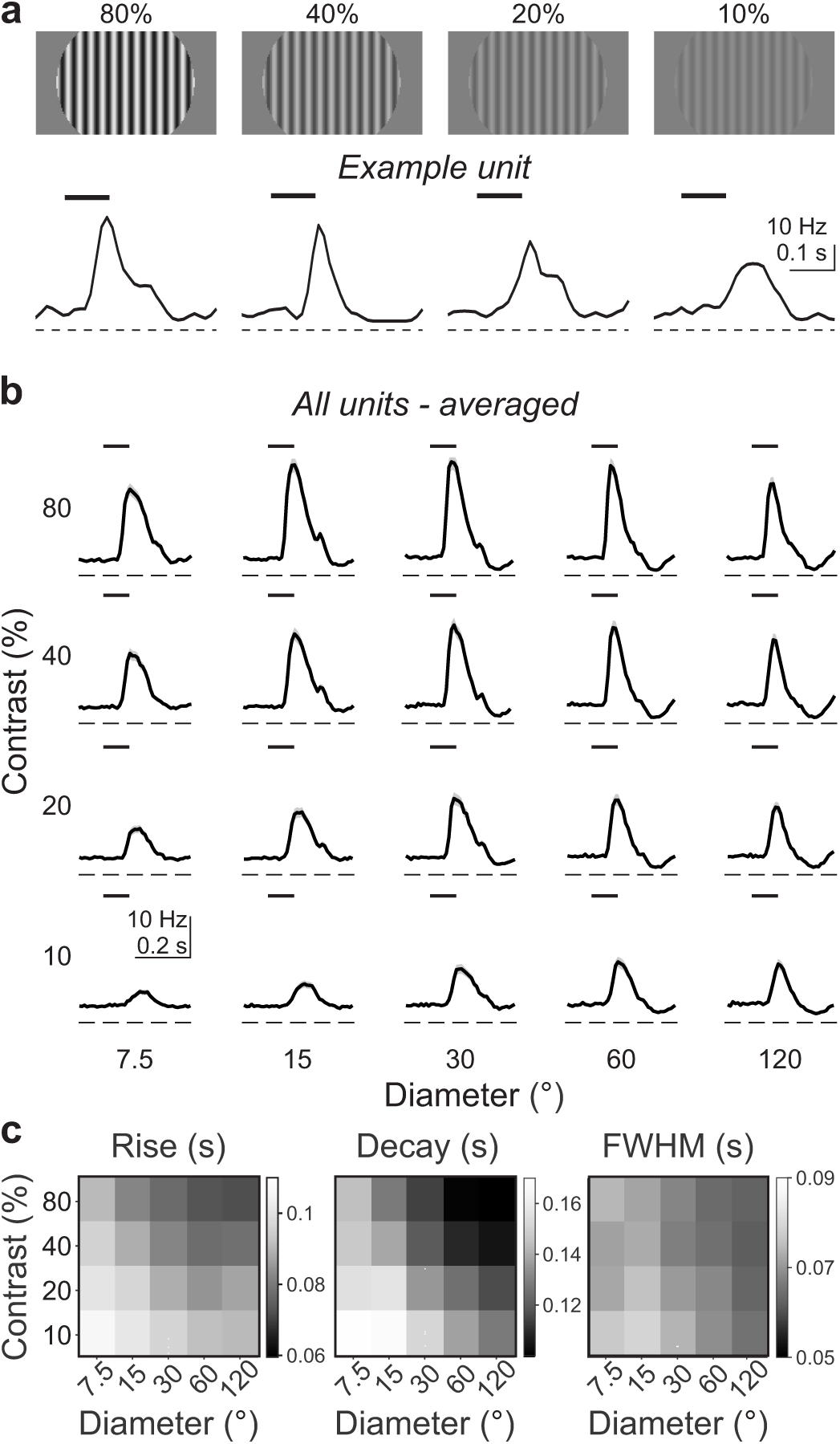
Increases in size and contrast differentially modulate response duration. (a) Visually-evoked PSTHs recorded in an example unit (same unit as in Figure 1g) during presentation of 120° diameter stimuli at different contrasts (10-80%). (b) Average visually-evoked PSTHs at all combinations of size and contrast (n = 321 units). Shaded error is SEM across units. (c) Left, heatmap of time to 50% rise, averaged across all units. Middle and right, same as left for 50% decay (middle) and FWHM (right).

To more directly test the contribution of size and contrast to visual response dynamics, we fit the rise, fall, and FWHM times with multiple linear regression models (**Methods**). Time to rise is more dependent on contrast than size (weights: contrast = −6.71e-03 ± 6.00e-04 (95% CI), size = −5.80e-03 ± 6.23e-04, p = 0.03, contrast vs size), whereas decay and FWHM are more dependent on stimulus size than contrast (decay weights: contrast = −8.34e-03 ± 9.46e-04, size = −1.23e2 ± 9.82e-04, p = 3.19e-09 contrast vs size; FWHM weights: contrast = −1.72e-03 ± 8.09e-04, size = −6.32e-03 ± 8.40e-04, p = 1.17e-15 contrast vs size). Thus, while contrast impacts the timing of visual responses and enhances the effects of surround suppression, size has a more robust effect on the decay, and subsequently the duration of visual responses.

To further investigate the effects of contrast on the dynamics of visual responses, we compared dPSTHs for increasing size at each contrast. The dPSTH for small sizes (e.g. comparing 7.5° vs 15°; **Figure 3a**) is primarily positive, independent of contrast. In comparison, the dPSTH for large size (i.e. comparing 7.5° vs 120°; **Figure 3b**) has a much weaker negative component at low contrast, rendering the integral of the dPSTH close to zero. As contrast increases, the integral goes significantly below zero, such that there is a significant dependence of the integral on contrast (two-way ANOVA, effect of contrast: p = 5.79e-34; interaction of size and contrast: p = 2.23e-09; **Figure 3c**). Thus, the decrease in firing rates are dependent on contrast, consistent with previous reports of surround suppression^35,36^.

**Figure 3.**
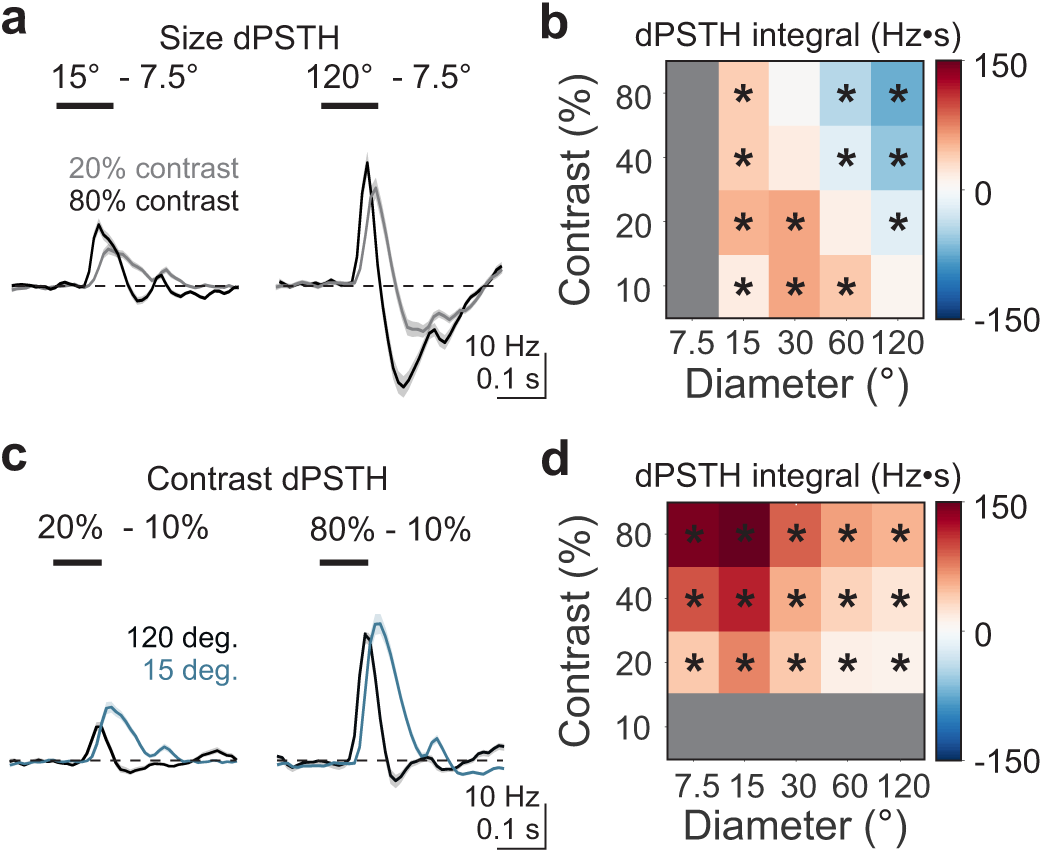
Increases in size uniquely alter dynamics of visually-evoked responses. (a) Average size dPSTH (comparing 7.5° and 15° stimuli [left] or 7.5° and 120° stimuli [right]) when the stimulus is 20% (light gray) or 80% (black) contrast (n = 321 units). Shaded error is SEM across units. (b) Heatmap of the dPSTH integral comparing across stimulus size; values significantly different from 0 (Student’s t-test) are noted with an asterisk. (c-d) Same as (a-b) for average contrast dPSTH, comparing 10% and 20% contrast (left) or 10% and 80% contrast (right) when the stimulus is 120°.

Compared to the dPSTHs for increasing size, the dPSTHs for increasing contrast have very different dynamics. The dPSTH for high contrast (i.e. comparing 20 vs 80% contrast) is monophasic and positive at both small and large sizes (**Figure 3d-e**). Indeed, the integral of the dPSTH between low and high contrasts remains positive, and becomes increasingly positive, with increasing size (two-way ANOVA, effect of contrast, p = 5.25e-42, interaction of contrast and size, p = 0.001; **Figure 3f**). As a result, whereas the integral of the dPSTH becomes smaller with increasing size, the integral becomes significantly larger with increasing contrast, reflecting a net increase in firing rate as a function of contrast. Thus, while the magnitude of surround suppression is modulated by contrast, the faster dynamics and lower firing rates that accompany surround suppression are exclusively evoked by increases in size.

### Visual response dynamics depend on a network mechanism

Surround suppression is thought to be engaged through the recruitment of lateral network interactions^17,29,37^. To determine whether the faster dynamics observed with increasing size could be explained by network mechanisms, or whether they are due to the recruitment of activity-dependent, cell-intrinsic mechanisms, we investigated the dependence of response dynamics on stimulus orientation. Given the robust orientation selectivity of V1 neurons^38^, if the observed dynamics reflect intrinsic properties, neurons should become more transient with increasing size only in response to their preferred orientation. Alternatively, if the observed dynamics reflect the broader recruitment of network mechanisms, then these phenomena should be independent of orientation.

Despite significantly weaker visual responses to the orthogonal stimulus (two-way ANOVA, effect of orientation: p = 7.42e-07; **Figure 4a-b**), there is still a robust dependence of average firing rate on stimulus size (effect of size: p = 5.84e-10). Visual responses continue to drive suppression of firing rates significantly below baseline following stimulus offset (**Figure 4a**). Indeed, for both preferred and orthogonal stimuli, size dPSTHs at large sizes are highly biphasic (**Figure 4c**), and the integrals of the size dPSTHs are significantly reduced with size (one-way ANOVA, effect of size, preferred: p = 1.23e-06, orthogonal: p = 3.02e-10). Given the smaller initial peak of the orthogonal stimulus, the integral of the orthogonal dPSTH is significantly more negative in response to the orthogonal stimulus (two-way ANOVA, effect of orientation, p = 1.12e-04; **Figure 4d**), but maintains the same size-dependent relationship (interaction of orientation and size, p = 0.77). Furthermore, we find no significant differences in the time to rise (two-way ANOVA, effect of orientation, p = 0.95; **Figure 4e**), decay (p = 0.25), or duration (p = 0.18) of visual responses. Thus, the strong suppression and faster dynamics are maintained in response to the orthogonal stimulus, consistent with the engagement of a network mechanism.

**Figure 4.**
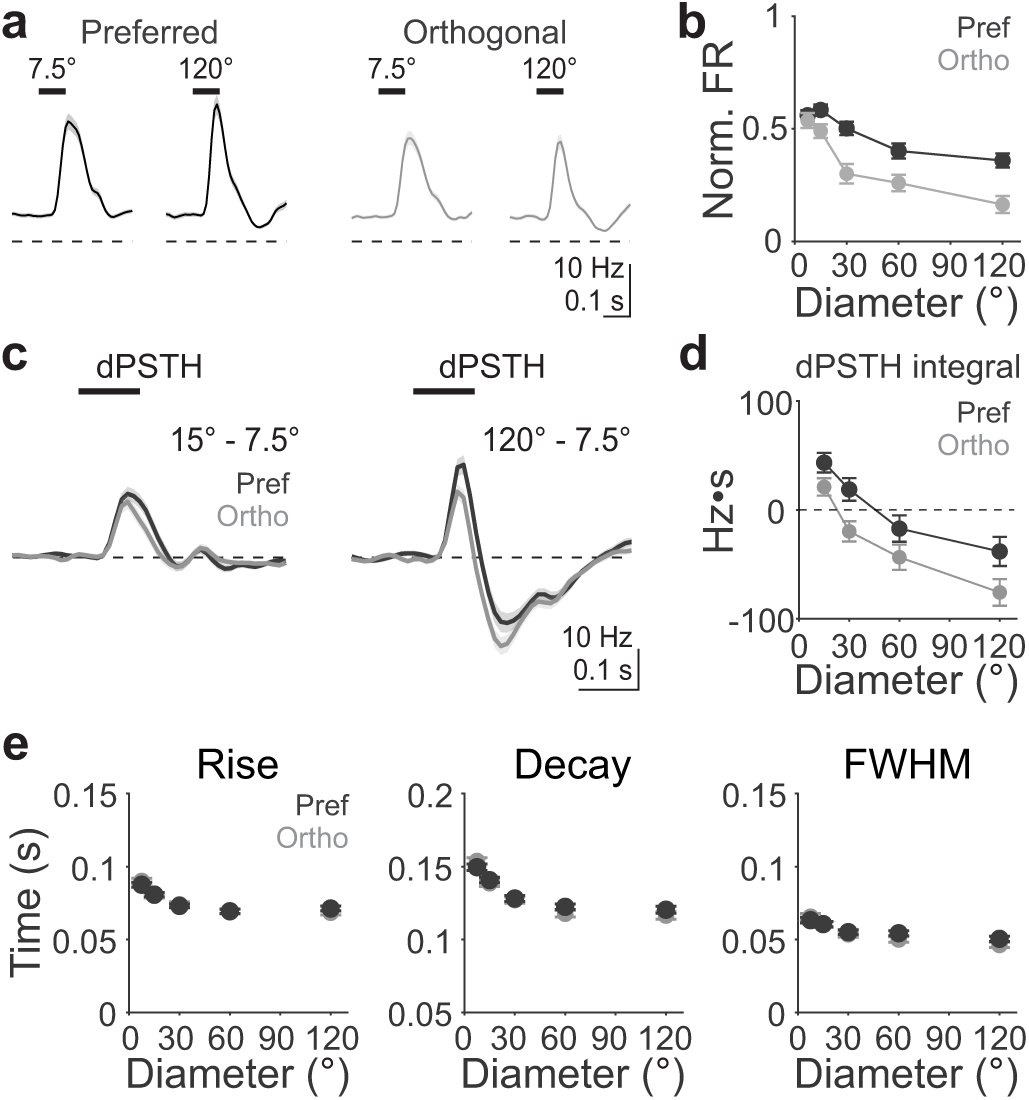
Size-dependent effects on response duration are independent of preferred orientation. (a) Average visually-evoked PSTH in response to presentation of preferred (left, dark gray) and orthogonal (right, light gray) gratings of 7.5° vs 120° diameter at 80% contrast (n = 321 units). Shaded error is SEM across units. (b) Average firing rate in response to preferred (dark gray) and orthogonal (light gray) stimuli as a function of stimulus size, normalized to the maximum response to the preferred stimulus. Error is SEM across units. (c) Left, dPSTHs for 7.5° vs 15° preferred (dark gray) and orthogonal (light gray) stimuli. Right, same as left for 7.5° vs 120° stimuli. (d) Average dPSTH integral in response to preferred (dark gray) and orthogonal (light gray) stimuli as a function of stimulus size. (e) Left, same as (d), for average time to 50% rise for cells responsive to each condition (n = 321, 289, 275, 252, 248). Middle and right, same as left for 50% decay (middle), and FWHM (right).

### Inhibitory cell-type specific dynamics with increasing size

The dependence of visual response dynamics on network mechanisms is consistent with current models of surround suppression that posit a role for the recruitment of local inhibition^9,29^. Somatostatin-expressing (SST) inhibitory interneurons respond preferentially to large, homogenous stimuli and have been specifically implicated in mediating surround suppression^17,30,39^. To determine whether SST cells also have size-dependent dynamics that could account for the dynamics of the responses in pyramidal cells, we performed multi-unit recordings in transgenic mice expressing the excitatory opsin channelrhodopsin-2 (ChR2) in SST cells in order to identify them through photo-tagging (n = 14 sessions, 9 mice; **Figure 5a**). All neurons that were not significantly driven were classified as either RS (putative pyramidal cells) or FS (putative parvalbumin-expressing (PV) cells) by the shape of their action potential waveform (**Figure S3a-b**).

**Figure 5.**
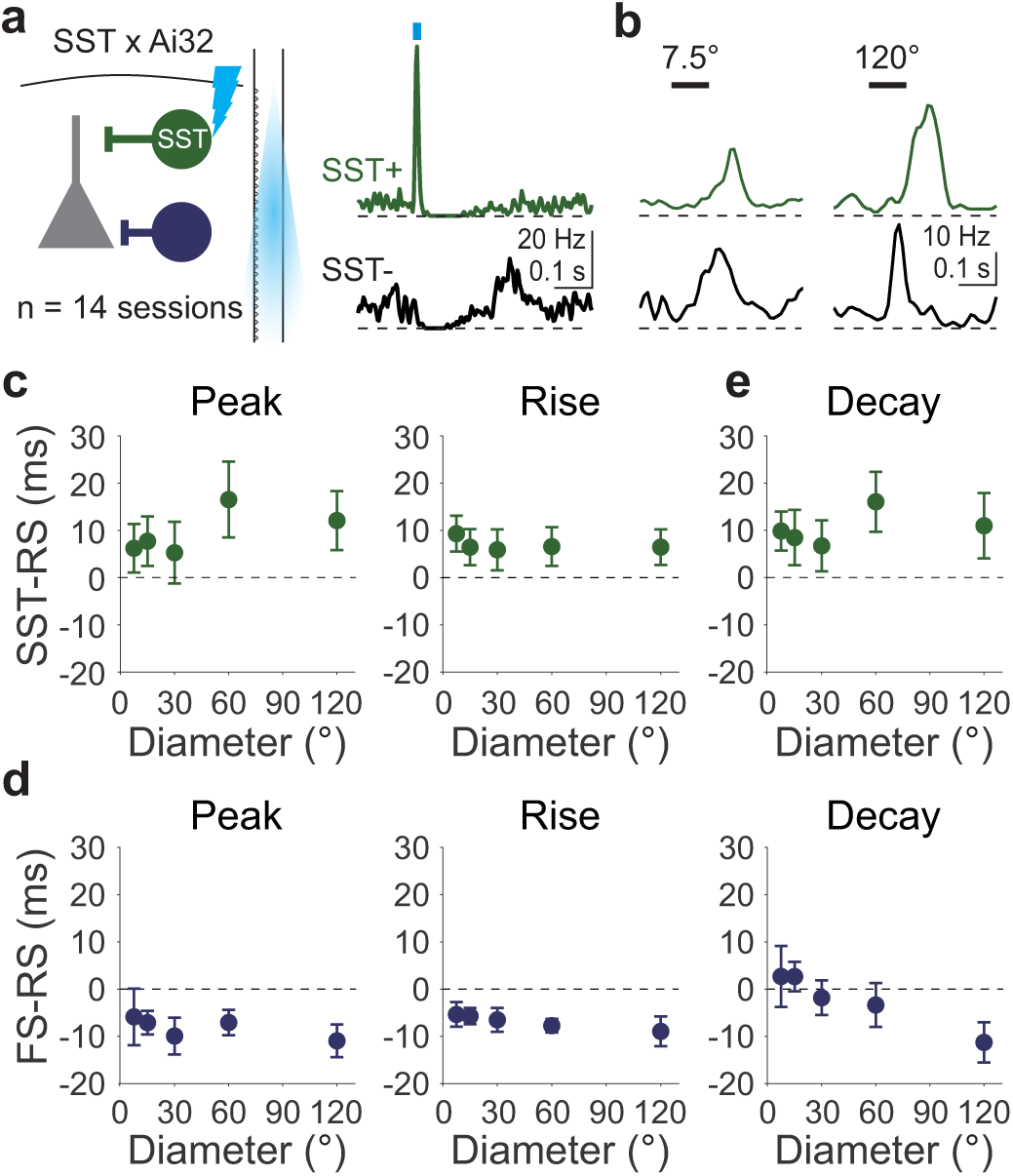
Somatostatin expressing (SST) cells have slower dynamics than regular-spiking (RS) and fast-spiking (FS) cells. (a) Left, schematic of recording configuration for opto-tagging SST cells in transgenic channelrhodopsin-2 reporter mice (Ai32). Right, example PSTHs at the time of optogenetic stimulation in an SST+ (top, green) and SST-(bottom, black) cell. (b) Example PSTHs in response to 7.5° (left) and 120° (right) stimuli for the same SST+ and SST-cells in (a). (c) Average difference in the time to peak (left), rise (middle) and decay (right) for SST cells and simultaneously recorded RS cells for sessions with at least one cell of each type responsive at that size (n = 12, 14, 14, 13, 13 sessions). (d) Same as (c), for FS cells (n = 14 sessions for all sizes).

We find that the selectivity and timing of visual responses of SST units (n = 56) are distinct from simultaneously recorded RS units (n = 351). SST cells have a larger preferred size (SST: 57.03 ± 6.56 deg; RS: 36.47 ± 2.30 deg; p = 0.01; unpaired t-test; **Figure 5b and S3c**) and the greatest number of SST neurons are significantly visually driven at the largest size (7.5 vs 120°: 42% vs 64%, p = 0.02; Chi-squared test; **Figure S3d**). Thus, consistent with previous work, SST cells are increasingly recruited with increasing stimulus size^17,37^. We next measured the average timing of visual responses in SST neurons relative to the simultaneously recorded RS units within each session. To limit our comparison to neurons with similar receptive field centers, we selected the subpopulation that was significantly responsive to the 15° stimulus (n = 31 SST units, n = 168 RS units). We find that the timing of visual responses in SST cells is delayed relative to RS neurons. SST cells have a later time to peak (repeated measures ANOVA, effect of cell type, p = 0.03; **Figure 5c**), rise (p = 0.04) and decay (p = 0.002), although the FWHM was not different (p = 0.16). Furthermore, the lag that SST cells exhibit in time to peak and decay are dependent on stimulus size (interaction of cell type and size-peak: p = 0.03; rise: p = 0.07, decay: p = 0.001). This suggests that a delayed, but not significantly prolonged, response in SST cells could truncate the response in RS cells to mediate the faster dynamics with increasing size.

To determine if these dynamics are specific to SST cells, we measured the dynamics of another population of putative inhibitory interneurons, the simultaneously recorded FS (n = 113) cells. In contrast to SST cells, preferred size of FS cells is not significantly different from RS cells (p = 0.81, un-paired t-test) and the greatest number of FS cells are activated at a smaller size, similar to RS cells (FS max 78% at 30 degrees; RS max 48% at 30 degrees; **Figure S3c-d**). The subset of retinotopically matched FS cells (n = 82 units, 14 sessions) exhibit significantly faster times to peak (repeated measures ANOVA, effect of cell type, p = 0.005; **Figure 5d**) and rise (p = 8.76e-5) compared to RS cells. Moreover, these differences are dependent on stimulus size (interaction of cell type and size-peak: p = 0.001; rise: p = 6.35e-5). In comparison, the time to decay and FWHM are not significantly different compared to RS cells (decay: p = 0.54; FWHM: p = 0.16). Thus, the selectivity and dynamics of visual responses in FS cells are not likely to account for the delayed suppression of visual responses in RS cells.

### Increasing stimulus size drives rapid suppression of both excitation and inhibition

The activation of SST cells may speed visual responses through increasing inhibition and hyperpolarizing their targets. Alternatively, activation of SST cells may trigger a broader network suppression where the initial hyperpolarization initiates a longer-lasting reduction in recurrent excitation^40^. In this model, network suppression reduces both excitation and inhibition with increasing stimulus size^29^. To understand how the underlying synaptic dynamics determine the strength and duration of visual responses, we made *in vivo* whole-cell recordings from neurons in L2/3 and measured synaptic input in response to stimuli of varying size (diameter: 7.5°, 30°, 120°; contrast: 80%; n = 14 cells, 7 mice). We voltage-clamped cells at −70 mV to record excitatory post-synaptic currents (EPSCs) and at +10 mV to isolate inhibitory post-synaptic currents (IPSCs; **Figure 6a-b**).

**Figure 6.**
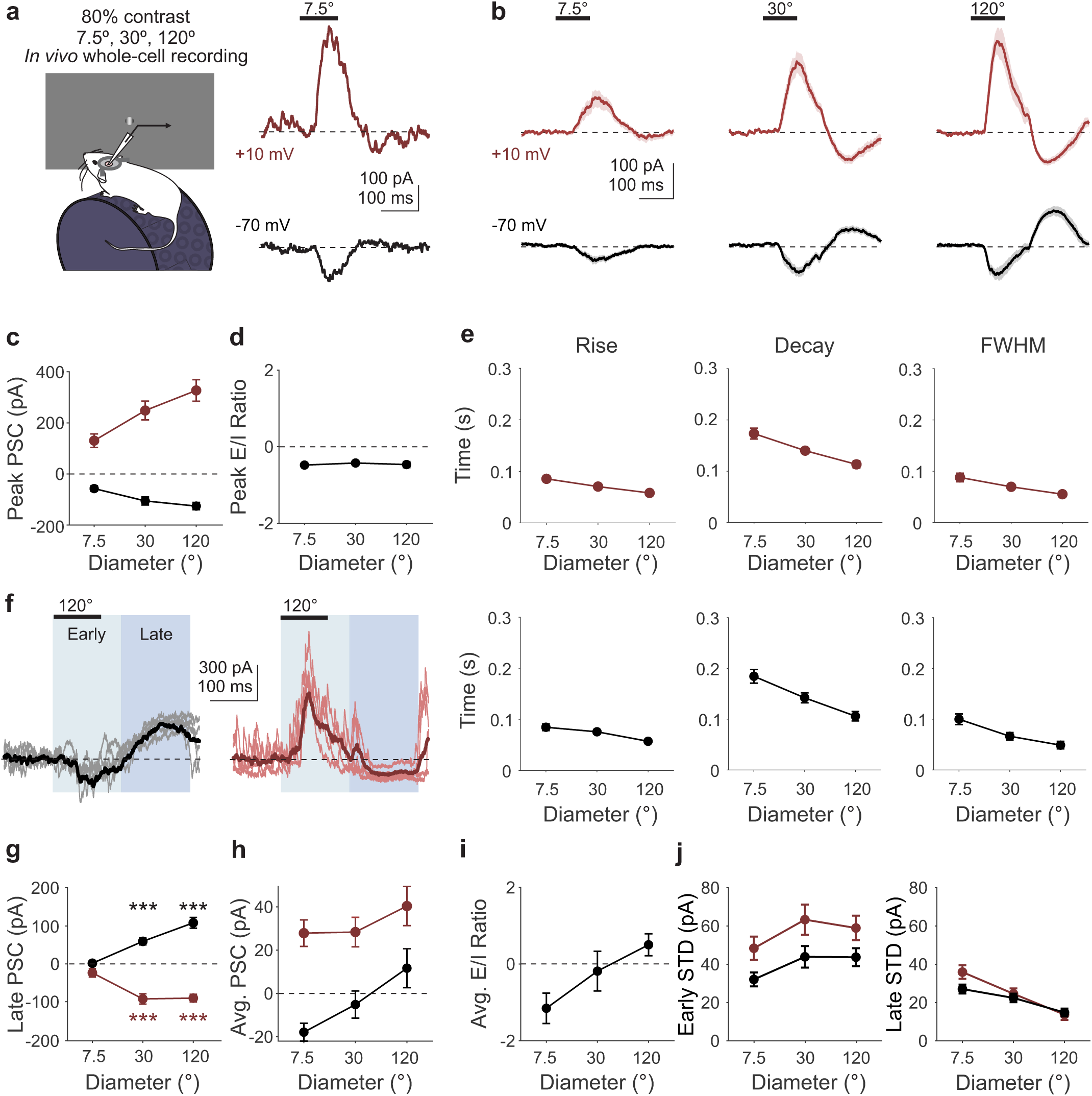
Network suppression underlies the size-dependent effects on response duration. (a) Left, schematic of intracellular recording configuration. Right, current traces from an example cell at the reversal potential for excitation (IPSCs; top, red) and inhibition (EPSCs; bottom, black) in response to a 7.5° stimulus. (b) Grand average IPSCs (top) and EPSCs (bottom) in response to stimuli of increasing size. Shaded error is SEM across cells (n = 13). (c) Average peak EPSC (black) and IPSC (red) amplitude as a function of stimulus size. Error is SEM across cells. (d) Average peak EPSC/IPSC (E/I) ratio as a function of stimulus size. (e) Time to rise (left), decay (middle) and FWHM (right) for IPSCs (top) and EPSCs (bottom) as a function of stimulus size for all cells (thick line) and individual cells (thin lines). (f) Example single trial traces (thin, light lines; n = 5) for EPSCs (left) and IPSCs (right) from a single cell. Average response (dark, thick line) is overlaid. Shaded regions identify early (light blue) and late (dark blue) analysis windows. (g) Same as (c), for PSC amplitude during the late analysis window. (h) Same as (c), for mean PSC amplitude. (i) Same as (d), for mean E/I ratio. Note that while E/I ratio appears to increase, this is due to a sign inversion where EPSCs become net positive, and therefore reflects a decrease in E/I ratio. (j) Same as (g), for the standard deviation (STD) of the early (left) and late (right) analysis windows.

Increasing stimulus size evokes EPSCs and IPSCs with increasing peak amplitude (repeated measures ANOVA, effect of size, p= 6.18e-05; **Figure 6c**). However, there is no significant dependence of the ratio of peak excitation to inhibition on stimulus size (E/I: one-way ANOVA, p = 0.79; **Figure 6d**). Thus, changes to the E/I ratio cannot explain the difference in firing rate or dynamics with increasing size.

Instead, as with the spike rates, the dynamics of both EPSCs and IPSCs become increasingly transient with stimulus size. Both EPSCs and IPSCs have faster rise (repeated measures ANOVA, effect of size, p = 1.25e-12, effect of current type, p = 0.71; **Figure 6e**) and decay (size, p = 3.45e-11, current type, p = 0.65) times with increasing size. For both EPSCs and IPSCs, the change in decay is larger than the change in rise, resulting in a shorter duration with increasing size (FWHM: effect of size, p = 3.04e-09, effect of current type, p = 0.84). Notably, there is no significant difference in the how dynamics of excitation and inhibition are modulated with size (interaction of current type and size, rise: p = 0.62; decay: p = 0.82; FWHM: p = 0.62). Thus, the increased transience of spiking with increasing size is not due to prolonged inhibition throughout the duration of suppressed activity.

Rather, we find that the faster dynamics in response to larger stimuli are due to a transient decrease in excitation below spontaneous levels (dark blue shaded region in **Figure 6f**; paired t-test with Bonferroni correction, late EPSC vs baseline, 7.5°: p = 1.00, 60°: p = 8.65e-05, 120°: p = 1.10e-05; **Figure 6g**), such that the average amplitude of the EPSC decreases with increasing size (one-way ANOVA, effect of size, p = 0.01; **Figure 6h**). Inhibition also decreases below baseline in the late period (paired t-test with Bonferroni correction, late IPSC vs baseline, 7.5°: p = 0.15, 60°: p = 4.08e-05, 120°: p = 2.17e-06; **Figure 6g**). However, the average IPSC decreases less with size than excitation (two-way ANOVA, interaction of size and current, p = 0.008; **Figure 6h**), resulting in a net shift towards inhibition and decrease in the E/I ratio (one-way ANOVA, p = 0.001; **Figure 6i**). The similarity of the timing of suppression of the EPSC and IPSC indicates that the decrease below baseline is unlikely to reflect voltage clamp errors (e.g. with IPSCs contaminating measures of EPSCs, or vice versa), and instead reflects a decrease in total input due to a network-wide decrease in excitability. Indeed, we find a significant decrease in the variance of both EPSCs and IPSCs during this late window (EPSC: p = 0.001, IPSC: p = 2.71e-05; one-way ANOVA; **Figure 6j**) consistent with a reduction in the total activity in the network. Therefore, the increased transience of spiking at large stimulus sizes arises from an increase in transience of both excitatory and inhibitory synaptic inputs due to a suppression of both below spontaneous levels.

### Locomotion reduces surround suppression by increasing response duration

Behavioral context has been demonstrated to broadly impact network state in cortex. Recent experiments reveal that locomotion makes visual responses more sustained^41^. In parallel, locomotion substantially relieves suppression^37,42,43^. To specifically test whether the effects of locomotion on surround suppression are due to a decrease in the transience of visual responses, we separated trials according to when the mouse was either stationary or moving during stimulus presentation. Notably, only a subset of experiments had sufficient locomotion trials for these analyses (n = 91 cells, 8 mice).

As with the full population, these units are significantly suppressed by large, high contrast stimuli when the animal is stationary (SI-0.62 ± 0.05; Student’s t-test, p = 1.09e-22). However, when animals are moving, we find that average visual responses are enhanced (repeated measures ANOVA, main effect of state, p = 0.003; **Figure 7a**) and less surround suppressed (SI-0.52 ± 0.06; interaction of state and size, p = 0.001), consistent with previous work^37,44,45^. With movement, there is no longer a strong suppression of firing rates below baseline in response to stimuli of large sizes (**Figure 7b**). These changes in the temporal dynamics of the response are reflected in the dPSTH which is more monophasic during locomotion than when stationary (**Figure 7c-d**). As a result, whereas the integral of the dPSTH in the stationary condition becomes significantly negative as size and contrast increase (similar to the full dataset, **Figure 3b**), none of the stimulus conditions have an integral significantly below zero when the mouse is running (**Figure 7e**), indicating a lack of suppression. Indeed, the integral is significantly increased during locomotion compared to the stationary condition across sizes (repeated measures ANOVA, effect of state, p = 0.03; **Figure 7e-f**). This is accompanied by a significant increase in the time to rise (repeated measures ANOVA, effect of state, p = 0.008; **Figure 7g**), decay (p = 1.12e-17), and duration (p = 3.98e-15) of visual responses during locomotion. Locomotion also reduces the size-dependence of dynamics across all measures (interaction of state and size, time to rise: p = 0.006; decay: p = 0.6.05e-17, FWHM: p = 7.07e-15). Thus, locomotion relieves surround suppression by prolonging the decay and preventing suppression below baseline firing rates for visually-evoked responses at large sizes.

**Figure 7.**
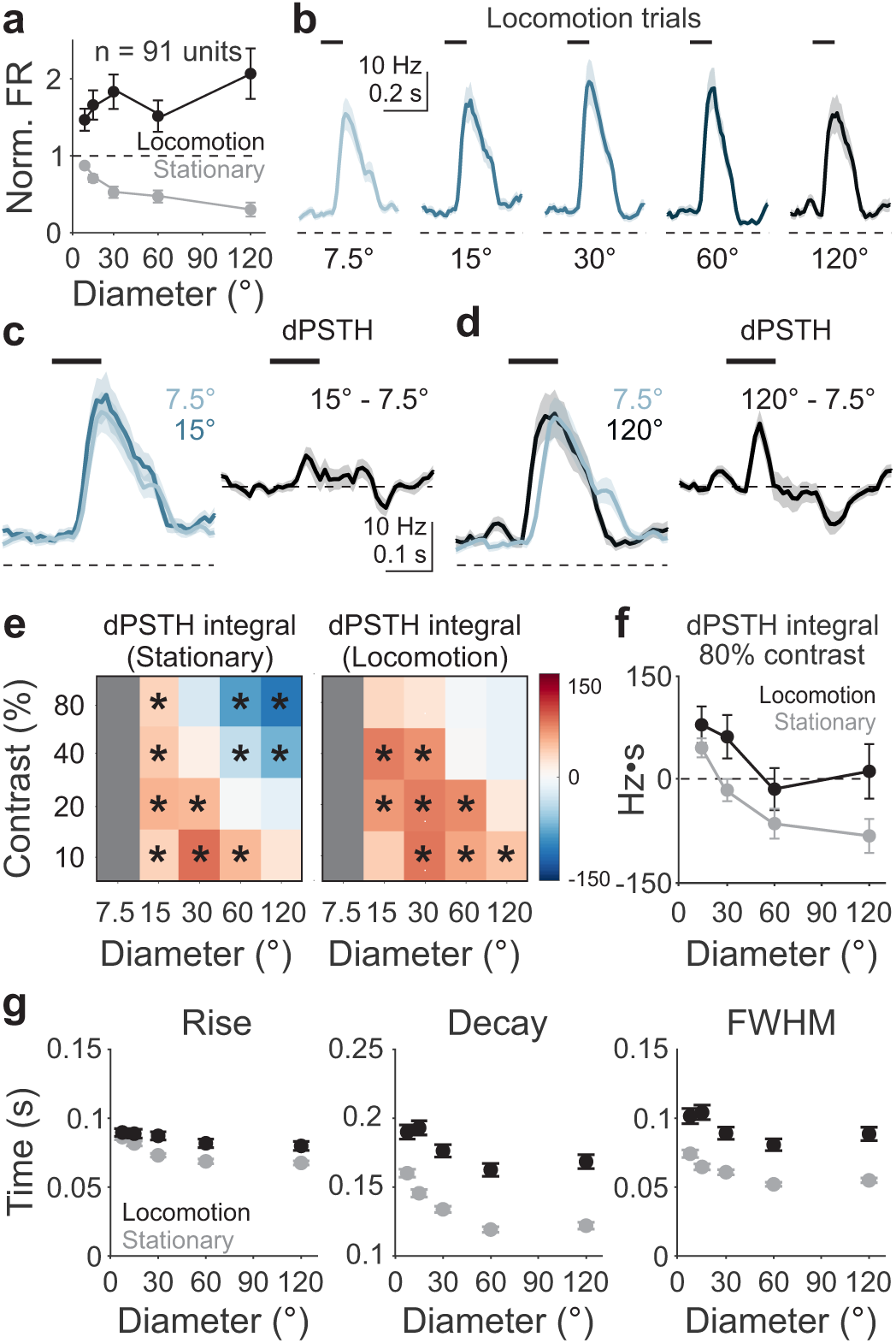
Locomotion counteracts size-dependent effects on response duration. (a) Average firing rate on stationary (gray) and locomotion (black) trials as a function of stimulus size at 80% contrast, normalized to the maximum response on stationary trials (n = 91 units). Error is SEM across units. (b) Average visually-evoked PSTH in response to presentation of static gratings of different diameters on locomotion trials. Shaded error is SEM across units. (c) Left, comparison of PSTHs for 7.5° (light blue) and 15° (dark blue) stimuli during locomotion; right, pointwise subtraction of stimuli on left are used to create the dPSTH. (d) Same as (c) but for comparison with responses to 120° stimulus. (e) Left, heatmap of the dPSTH integral comparing across stimulus size for stationary trials; values significantly different from 0 (Student’s t-test with Bonferroni correction) are noted with an asterisk. Right, same as left, for locomotion trials. (f) Same as (a), for average dPSTH integral at 80% contrast. (g) Left, same as (a), for average time to 50% rise for cells responsive to each condition (n = 91, 82, 73, 68, 65). Middle and right, same as left for 50% decay (middle), and FWHM (right).

## Discussion

Here we show that the network mechanisms that regulate spatial integration in mouse V1 do so by altering temporal dynamics of visually-evoked responses. Increasing stimulus size transiently recruits strong and balanced synaptic excitation and inhibition; however, following this transient increase in excitation and inhibition, network activation is rapidly suppressed, resulting in a decrease in both excitation and inhibition below baseline levels. The rapid synaptic dynamics evoked by large stimuli decrease the duration of visual responses, leading to the decreased firing rates observed during surround suppression.

While both surround suppression and size-dependent changes in response dynamics are present in the retina and lateral geniculate nucleus (LGN), we think that the majority of our observations are due to cortical dynamics. First, robust suppression of peak firing rates is observed in the retina and LGN^15^, whereas we find relatively weak effects on the peak compared to the average firing rate in V1. Second, we find that stimulus size has an increasing effect on both firing rates and decay time out to the largest size presented (120°). This is well beyond the range over which pre-cortical circuits are thought to integrate^13,15,16^. Third, the effect of size on temporal integration in the retina and LGN is considered to be a form of contrast adaptation, as similar effects are generated by increasing contrast. However, we find that the effects of contrast and size can be dissociated in the cortex, suggesting that they have distinct underlying mechanisms. The effects of contrast are thought to be partly inherited from the LGN and amplified by cortical short-term depression^46^. We propose that the effect of size is largely due to the intrinsic dynamics of the cortical circuit.

We were able to reveal how stimulus features influence network dynamics by presenting brief, stationary stimuli. Prolonged stimuli evoke adaptation and motion drives complex feedforward dynamics, which are difficult to disentangle from the intrinsic dynamics of the cortical network. Using this brief stimulus, we could transiently activate cortex and such that the cortical PSTH reveals the inherent temporal characteristics of the system, akin to an impulse response function. We find that this impulse response function depends on both stimulus contrast and size. Whereas increases in contrast and size both speed the time to rise and decay of visual responses, increasing size has a disproportionate effect on the decay time, resulting in a selective decrease of the duration of visual responses. This is due to the biphasic nature of the impulse response function at large sizes, where both firing rates and synaptic currents are only transiently increased before being driven below baseline.

The dynamics that we observe are consistent with predictions of recurrent networks. When recurrent excitation is strong enough to require recurrent inhibition for stability (i.e. the cortex is an ISN), the network undergoes unique temporal dynamics: strong excitation recruits robust recurrent inhibition, which in turn suppresses local excitation and thus removes excitatory drive from both excitatory and inhibitory populations^27,28^. Thus, sufficiently strong input can drive a paradoxical suppression of the network. This network suppression has been previously proposed to underlie surround suppression^29^, and has also been observed in response to optogenetic stimulation of visual cortex^31,47^ and presentation of non-preferred tones in auditory cortex^40^. Here we demonstrate temporal dynamics of network suppression that are dependent on contrast and independent of orientation preference, which are altogether consistent with the engagement of an ISN.

Notably, we find no size-dependence of the excitation to inhibition (E/I) ratio of the peak response. This is contrary to previous experiments measuring E/I ratios in response to drifting gratings^30^. Those experiments revealed that excitation and inhibition both increase with stimulus size, but that inhibition increases more than excitation, thereby decreasing the E/I ratio. We do find a small decrease in the E/I ratio of the average response, but this arises from a stronger suppression of excitation below resting levels, rather than a relative increase in inhibition. In addition to the different stimulus duration and motion, there are a couple of other differences between our experiments that could explain the discrepancies in the effect on E/I ratio. First, we did not include the sodium channel blocker QX-314 in our internal solution as this has been shown to reduce the degree of spontaneous activity, and therefore the magnitude of network suppression^40^. Second, the majority of our data was collected in the stationary condition, whereas the finding of decreased E/I ratio was exclusively during locomotion.

Indeed, spatial and temporal integration in the visual system are sensitive to behavioral state. Locomotion both reduces surround suppression^37,42^ and prolongs visual responses^41^. Our data suggest that these effects may arise from a shared underlying mechanism that modulates response duration. We hypothesize that locomotion reduces the recruitment of synaptic inhibition (likely by reducing synaptic output as the firing rates of most interneurons increase^37,45,48^), relieving the strong network suppression that pushes firing rates below spontaneous levels. Unfortunately, we were not able to collect sufficient trials during locomotion in our intracellular recordings to test whether locomotion in fact reduces the suppression of synaptic input. However, comparable recordings from the auditory cortex during locomotion find that network suppression is reduced during locomotion^49^. Thus, these data provide a mechanistic account for how behavioral state regulates sensory integration.

SST-expressing interneurons have been proposed to drive the inhibition underlying surround suppression^17^. Indeed, we find that the pattern of recruitment of SST cells is consistent with this role. Consistent with previous studies, we find that SST cells are more strongly recruited with increasing stimulus size^17,37^. Moreover, we find that the recruitment of SST cells is delayed relative to simultaneously recorded pyramidal cells, especially at larger sizes. These slower dynamics could support their role in providing recurrent inhibition to their neighbors and initiating network suppression. Notably, although SST cells exhibit delayed dynamics, they are also ultimately suppressed below baseline levels, consistent with their engagement in the ISN (**Figure S3**). Given the prolonged inhibition provided by SST cells, we might have expected to see a transient decrease in the E/I ratio. It is possible that we may not detect the extra inhibitory input due to the distal location of SST synapses. In our somatic recordings, the inhibitory currents are likely dominated by the faster dynamics of inputs from FS (putative PV) cells. Nonetheless, our recordings likely reflect the summed input that influences spiking output, and therefore serve as a good proxy for pyramidal cell excitability.

In summary, using *in vivo* recordings of neurons in mouse primary visual cortex, we have linked the widely observed phenomenon of surround suppression with temporal dynamics of neural responses that are dependent on cortical network state. We find evidence for a mode of cortical function in which increasing stimulus size drives stabilizing network suppression, which is reflected in an acceleration of the dynamics of neural responses and decreasing average firing rates. Altogether, these data present a mechanistic link between spatial and temporal integration in the visual system that is likely to be conserved across brain areas^40^ and species^29^, and open up future investigations into how different stimulus features are integrated in cortical networks across behavioral states.

## Data availability

All data and code needed to reconstruct the figures is available on figShare (doi: 10.6084/m9.figshare.28711241). Requests for access to raw datasets can be made to the corresponding author at.

## Acknowledgements

We thank Wenjuan Kong and Gloria Kim for assistance with husbandry, surgeries and retinotopic mapping. We thank Dr. Nicolas Brunel for comments on the manuscript and members of the Hull and Glickfeld labs for insight throughout the project. This work was supported by grants from the National Institutes of Health (R01-EY031716 to L.L.G., F31-EY031941 to J.Y.L., and F32-EY034013 to C.M.C.).

## Author contributions

Conceptualization: J.Y.L., C.M.C. and L.L.G; Investigation and data curation: J.Y.L.; Formal analysis: J.Y.L.; Writing-Original draft: J.Y.L. and L.L.G; Writing-Review and editing: J.Y.L., C.M.C. and L.L.G; Supervision: L.L.G.; Funding acquisition: J.Y.L., C.M.C. and L.L.G.

## Declaration of interests

The authors declare no competing interests.

## Figure Legends

**Supplementary Figure 1.**
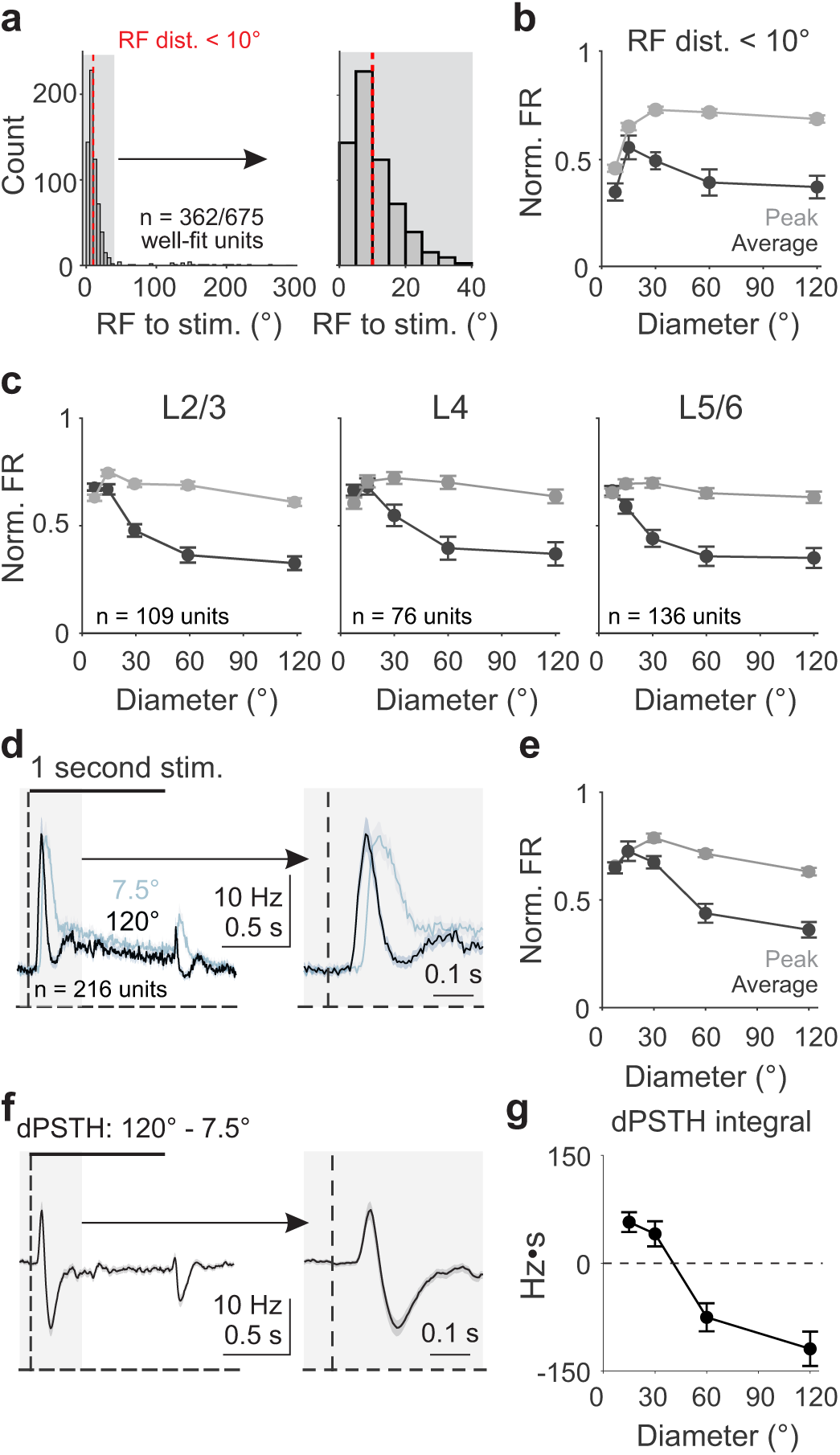
Size-dependent visual response dynamics are not dependent on unit selection method, cortical layer or stimulus duration, related to Figure 1. (a) Left, distribution of distance from the stimulus to receptive field center for all well-fit units (n = 675 units). Vertical red line delineates units within 10 degrees of the stimulus (n = 362 units). Right, expansion of shaded region on left. (b) Average normalized firing rate for units within 10 degrees of the stimulus as a function of stimulus diameter measured using the average (dark grey) and peak (light grey) of the PSTH. Error is SEM across units. (c) Same as b, for units in L2/3 (left), L4 (middle) and L5/6 (right). (d) Left, average PSTH in response to presentation of 7.5° (light blue) and 120° (black) static stimulus for 1 s (n = 216 units). Shaded error is SEM across units. Black bar is stimulus duration. Right, expansion of shaded region on left. (e) Same as (b), for 1 s stimulus. (f) Left, dPSTH for comparison of 7.5° and 120° 1 s stimuli. Right, expansion of shaded region on left. (g) Integral of the dPSTH for 1 s stimuli as a function of stimulus diameter.

**Supplementary Figure 2.**
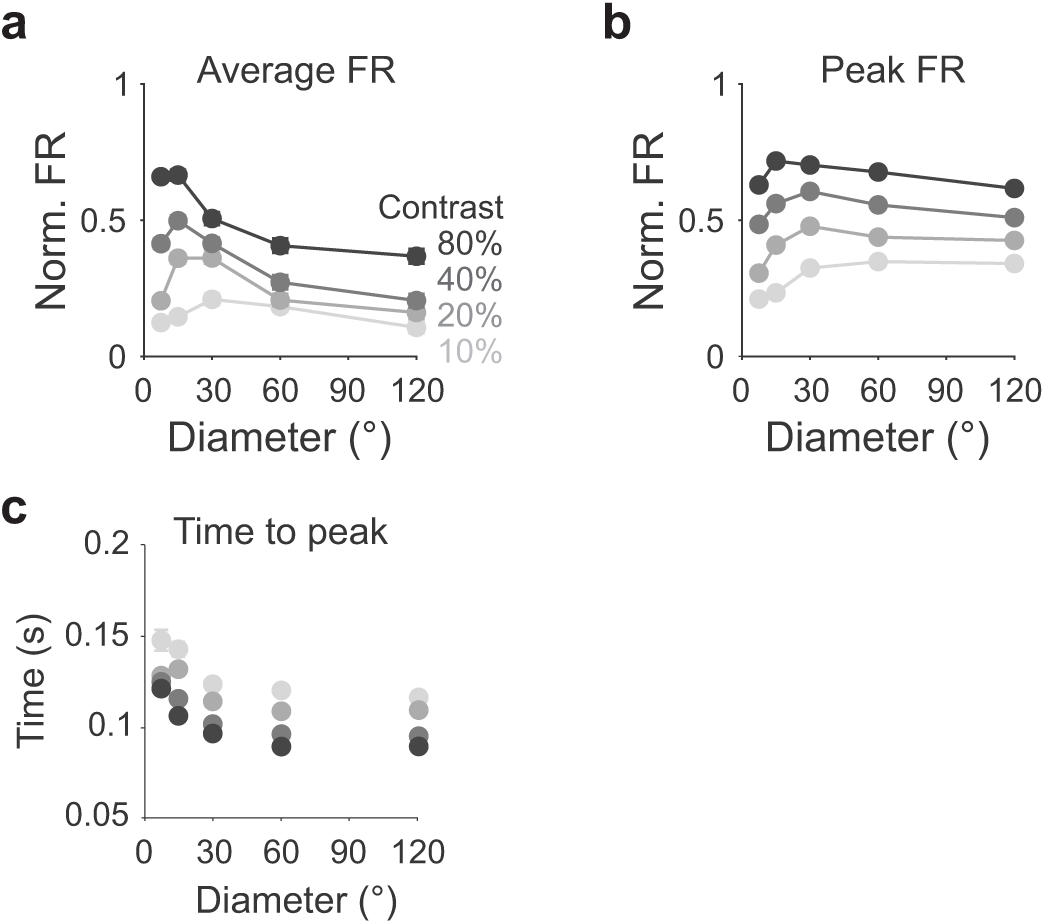
Size-dependent visual response dynamics are modulated by stimulus contrast, related to Figure 2. (a) Average normalized firing rate as a function of stimulus diameter and contrast (shades of gray) measured using average of the PSTH. Error is SEM across units. Two-way ANOVA, interaction of size and contrast: p = 2.04e-08. (b) Same as (a), measured using the peak of the PSTH; p = 0.10. (c) Same as (a), for time to peak; p = 0.02.

**Supplementary Figure 3.**
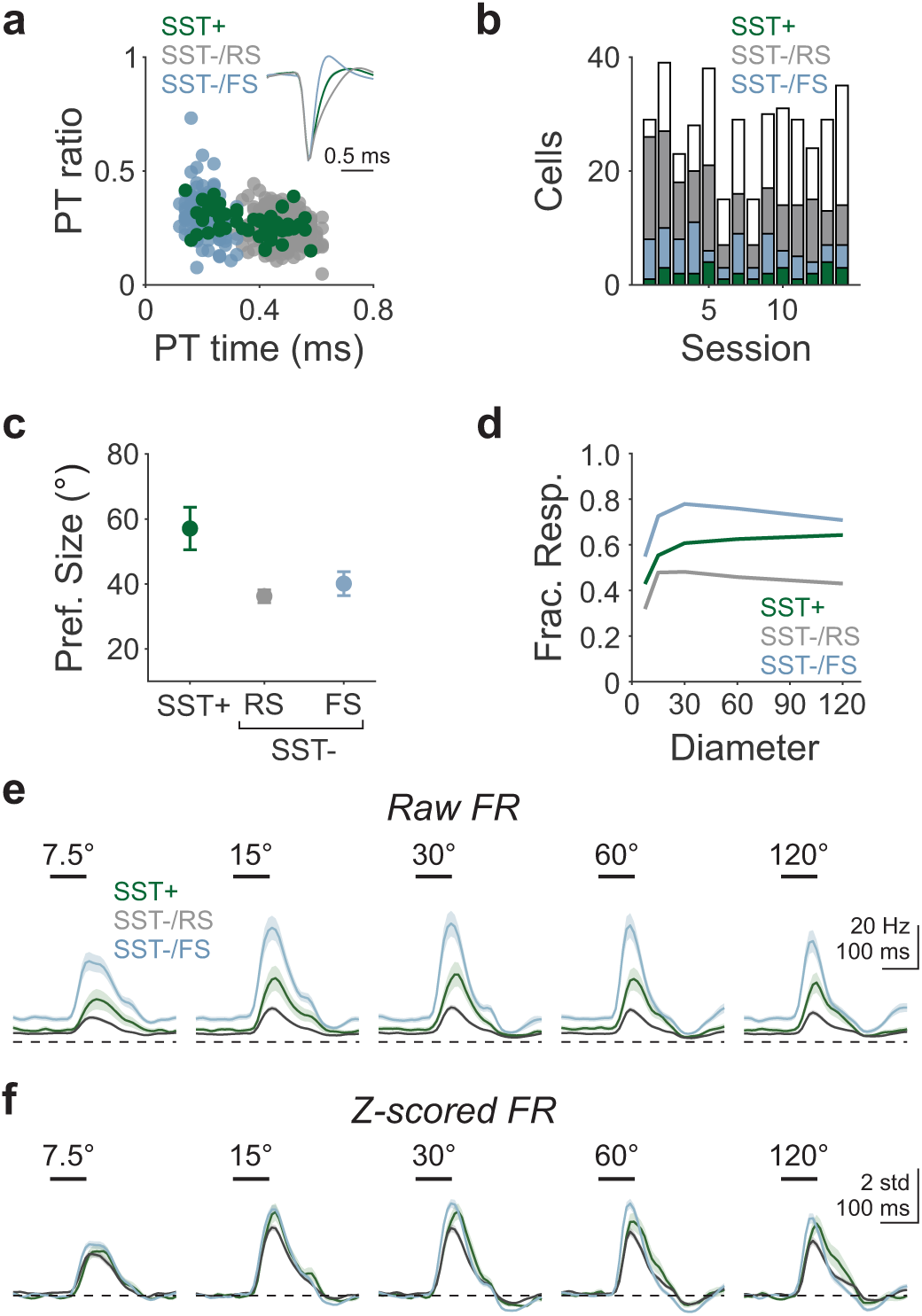
Identification and comparison of SST, regular spiking (RS) and fast spiking (FS) cells, related to Figure 5. (a) Scatter of peak-to-trough (PT) time versus ratio for all optogenetically identified SST+ (green; n = 56), and non-tagged SST-/RS (gray; n = 351) and SST-/FS (light blue; n = 113) cells. Inset, average spike waveform for the three classes of neurons. Shaded error is SEM across cells. (b) Number of SST+ (green), PV-/RS (gray), PV-/FS (light blue), and non-visually driven (white) cells in each recording session (n = 14 sessions, 9 mice). (c) Average preferred size for the cells in (b). Error is SEM across cells. (d) Fraction responsive as a function of stimulus size for the cells in (b). (e) Average PSTH of units across stimulus size, selecting units visually responsive to the 15° stimulus. (f) Z-scored PSTHs for units in (e).

## Methods

### Experimental Methods

#### Animals

All procedures conformed to standards set forth by the National Institutes of Health Guide for the Care and Use of Laboratory Animals, and were approved by the Duke University’s Animal Care and Use Committee. Mice were housed on a normal 12:12 light-dark cycle. Data in this study were collected from 38 mice (14 female). Experiments involving optogenetic tagging of SST interneurons used offspring from SST-Cre mice (Jackson Labs #013044) crossed with Ai32 (Jackson Labs #012569, n = 9). All other experiments did not require cell-type specific expression; thus, mice were a mix of genotypes. Transgenic mice were heterozygous and bred on a C57/B6J background (Jackson Labs #000664). Animals were 8-20 weeks old at the time of recording.

#### Headpost implantation

Mice were anesthetized with a mixture of ketamine/xylazine (ketamine: 50 mg/kg, xylazine: 5 mg/kg; intraperitoneal) and isoflurane (1.2–2% in 100% O_2_). Meloxicam was administered pre-operatively (1 mg/mL, 5 mg/kg; subcutaneous). Using aseptic technique, a custom-made titanium headpost was secured over V1 using clear dental cement (C&B Metabond, Parkell). Buprenex (0.05 mg/kg) and cefazolin (50 mg/kg) were administered post-operatively. Animals were allowed to recover for at least 1 week prior to experiments.

#### Visual stimulus presentation

Visual stimuli were presented on a 144-Hz (Asus) LCD monitor, calibrated with an i1 Display Pro (X-rite). The monitor was positioned 21 cm from the contralateral eye. Visual stimuli were controlled with MWorks (http://mworks-project.org). To measure size and contrast tuning, circular gabor patches containing sine-wave gratings (7.5°-120° diameter; 0.1 cycles per degree; 10%-80% contrast) alternated with periods of uniform mean luminance (60 cd/m^2^). Gratings were randomized in orientation (0°, 45°, 90°, 135°). For receptive field mapping, we presented flashing horizontal and vertical white bars (5° width). Timing of visual stimulus onset was measured via a photodiode that directly measured output from the LCD and used to align neural data.

#### In vivo retinotopic mapping

For all *in vivo* electrophysiological recordings, V1 boundaries were first identified with retinotopic mapping with intrinsic signal imaging through the skull. The skull was illuminated with orange light (590 nm LED, Thorlabs), and unfiltered emitted light was collected using a CCD camera (Rolera EMC-2, Q Imaging) at 2 Hz through a 5x air immersion objective (0.14 numerical aperture (NA), Mitutoyo), using Micromanager (NIH) acquisition software. Drifting gratings (80% contrast, 2 Hz, 0.1 cpd) were presented for 2 s at 3 positions with a 4 s interstimulus interval. Collected images were analyzed in ImageJ (NIH) to measure changes in reflectance at each position (dR/R; with R being the average of all frames) to identify V1.

#### Preparation for in vivo electrophysiology

Animals were habituated to head-fixation for 1-3 days prior to surgery. The day of recording, animals were anesthetized with isoflurane and a small craniotomy (< 1 mm diameter) was made over a V1 location identified by intrinsic signal imaging. For extracellular recordings, a gold ground pin was inserted in an anterior portion (outside of visual areas) within the headpost and secured with dental cement. Damage to superficial cortex was minimized by drilling in brief bouts (< 1 s) and alternating drilling and cooling with chilled glucose-free HEPES-based ACSF (in mM: 141 NaCl, 2.5 KCl, 10 HEPES, 2 CaCl_2_ 1.3 MgCl_2_). A slit was made in the dura with a syringe and the craniotomy was kept covered with ACSF for the remainder of the experiment. Animals were allowed to recover on the running wheel for at least 45 minutes before recording. In a subset of experiments, recording was performed the day after the craniotomy or animals were used for up to 2 consecutive recording days. In these cases, the craniotomy was protected overnight with Dura-Gel (Cambridge NeuroTech) and dental cement, which were removed and replaced with ACSF prior to recording.

#### In vivo whole-cell recordings

Whole-cell recordings were performed using blind patch technique. A silver chloride ground pellet was placed in the recording well outside of the brain. Recording ACSF was wicked away from the craniotomy and a 3-5 MOhm glass micropipette was lowered until the pipette tip touched the brain (confirmed by appearance of a square pulse on the membrane test); this position was zeroed and the well was refilled with recording ACSF. All recordings were documented relative to this depth. The pipette was lowered to ∼100 μm depth and then stepped in 1-2 μm increments until an increase in resistance was observed, at which point pressure was released to form a GΩ seal. Cells recorded at 180-350 μm depths were considered to be within L2/3. For voltage clamp recordings, internal solution contained (in mM): 125 Cs-methanesulfonate, 5 TEA-Cl, 10 HEPES, 0.5 EGTA, 4 Mg-ATP, 0.3 Na_3_GTP, 8 phosphocreatine-di(tris), 3 NaCl. EPSCs were recorded at −70 mV and IPSCs were recorded at +10 mV based on our calibration with ChR2 activation of interneurons *in vivo*^50^. Series resistance was monitored using −5 mV steps preceding each stimulus; recordings that reached >35 MΩ resistance or >20% change from baseline were discarded.

#### In vivo extracellular recordings

At the time of recording, animals were head-fixed and placed on a custom-built foam treadmill. Extracellular recordings were performed with a 32-site silicon probe (A1×32-Poly2–5mm-50s-177-A32, NeuroNexus or H4, Cambridge NeuroTech). Probes were connected through an A32-OM32 adapter to a Cereplex Mu digital headstage (Blackrock Microsystems). Signals were digitized at 30 kHz and recorded by a Cerebus multichannel data acquisition system (Blackrock Microsystems). Probes were slowly lowered into the brain until all sites were inserted and allowed to stabilize for 40–50 min before recording. Locomotion data was recorded using a rotary encoder built into the treadmill.

For optogenetic tagging experiments, we used a 450 nm laser (Optoengine) coupled to an optic shutter and patch cable terminating in an optic fiber. Optic fibers were either flat and terminated 100 μm above the surface of the brain (200 μm core, 0.22 NA), or were a tapered lambda fiber (100 μm core with 0.9 mm taper, 0.22 NA, Optogenix) aligned to the tip of the probe for enhanced light transmission deeper in the brain. Laser power was calibrated to deliver 2 mW power at fiber tip for ChR2 activation. Putative ChR2-expressing units were identified by significant, short-latency (< 3 ms) increase in firing in response to ∼50 pulses of light (10 ms, 0.1 Hz).

### Analysis of extracellular recordings

#### Spike sorting

Single units were isolated with KiloSort 2.5 (https://github.com/MouseLand/Kilosort) using refractory period violations and steepness of the autocorrelogram as criteria for isolation. We then manually curated these units in Phy (https://github.com/cortex-lab/phy) such that only units that were detected throughout the entire recording were included for subsequent analysis. Depth of the unit was assigned based on their waveforms’ center-of-mass across sites. Fast-spiking (FS) and regular-spiking (RS) units were separated within recordings according to peak-to-trough time of the maximum amplitude waveform across all contact sites (**Figure S3**).

#### Layer identification

To functionally identify cortical layers, we used the local field potential (LFP) obtained from filtering the raw data (downsampled to 10 kHz) from 1 to 200 Hz. The trial-averaged, stimulus-evoked LFP during a 1-second drifting grating presentation was converted to a current source density (CSD) plot by taking the discrete second derivative across the electrode sites and interpolated. Layers bounds were assigned relative to an initial sink in layer 4, followed by a sink in layer 2/3 and a sustained sink in layer 5.

#### Identification of locomotion trials

In all figures except **Figure 7**, only stationary trials were used. To categorize stimulus presentations as occurring during stationary or locomotion, digital signal from the encoder was converted to running speed using the diameter of the circular treadmill. If the average running speed in a 0.5 s window starting 0.4 s before stimulus onset until stimulus offset exceeded 10 cm/s, the trial was categorized as a locomotion trial.

#### Quantification and Statistical Analysis

All analyses were performed in custom code written in either MATLAB or Python. All data are presented as mean ± SEM. N values refer to number of cells or units isolated. Sample sizes were not predetermined but are comparable to published literature for each type of experiment. All statistics are corrected for multiple comparisons.

#### Data inclusion and analysis

For all extracellular recordings, PSTHs were generated by binning spiking activity in 0.01 s windows across all trials of each type, aligned to stimulus onset. For all figures except **Figure 4**, responses were collapsed across orientation. Visual responsiveness within each stimulus condition was measured using a paired t-test in a 0.15 s window before and after stimulus onset for trials of that type. For extracellular recordings, the peak response was measured as the maximum firing rate from 0-0.35 s after stimulus onset. For intracellular recordings, peak current was measured as the mean current in a 0.05 s window around the minimum (EPSC) or maximum (IPSC) value. The maximum late suppression was measured as the mean current in a 0.05 s window around the maximum (EPSC) or minimum (IPSC) value 0.15-0.3 s after stimulus onset. For both extracellular and intracellular recordings, the average response was measured as the mean current or firing rate in the full 0-0.35 s time window. For all measurements, we performed a baseline subtraction using a 0.15 s baseline window prior to stimulus onset.

Suppression index (SI) was calculated as the ratio of the response to the largest size and the response to the preferred size. Delta PSTHs (dPSTHs) were calculated across sizes by subtracting off the response to the smallest size (7.5°) or across contrasts by subtracting off the response to the smallest contrast (10%). dPSTH integral was calculated as the sum of the dPSTH 0-0.35 s after stimulus onset.

To measure peak, rise, decay, and full-width half-max (FWHM) in extracellular recordings, visually-evoked responses 0-0.35 s after stimulus onset were smoothed with a 50 ms gaussian window, and then resampled with interpolation at 10 times the original binning (yielding 0.001 s bins). For intracellular recordings, rise, decay, and FWHM measurements were made using the smoothed (10 ms gaussian), trial-averaged current response at each size. We measured the peak as the maximum response, the rise as the first time the response exceeded 50% of the maximum response, the decay as the first time the response dropped below 50% of the maximum response, and the FWHM as the time between these two points. To determine the relative influence of size and contrast on rise, fall, and FWHM, we performed a linear regression using z-scored size and contrast as predictor variables and FWHM duration as the response variable to calculate the coefficient (slope) of each of these features.

To measure the fluctuations of intracellularly recorded currents as a proxy for spontaneous activity levels, we measured the standard deviation of the currents on single trials in a 0.15 s window starting immediately after stimulus onset (Early; 0-0.15 s) or 0.15 s after stimulus onset (Late; 0.15-0.3 s).

## Notes

### Competing Interest Statement

The authors have declared no competing interest.

